# Serum iron variation is circadian-regulated and linked to the harmonic circadian oscillations of erythropoiesis and hepatic *Tfrc* expression in mice

**DOI:** 10.1101/2023.05.07.539729

**Authors:** Cavan Bennett, Anne Pettikiriarachchi, Alistair R.D. McLean, Rebecca Harding, Marnie E. Blewitt, Cyril Seillet, Sant-Rayn Pasricha

**Affiliations:** Population Health and Immunity Division, The Walter and Eliza Hall Institute of Medical Research, Parkville, Victoria, Australia; Department of Medical Biology, University of Melbourne, Parkville, Victoria, Australia; Centre for Epidemiology and Biostatistics, Melbourne School of Population and Global Health, The University of Melbourne, Melbourne, Australia; Epigenetics and Development Division, The Walter and Eliza Hall Institute of Medical Research, Parkville, Victoria, Australia; Immunology Division, The Walter and Eliza Hall Institute of Medical Research, Parkville, Victoria, Australia; Diagnostic Haematology, The Royal Melbourne Hospital, Parkville, Victoria, Australia; Clinical Haematology at the Peter MacCallum Cancer Centre and The Royal Melbourne Hospital, Melbourne, Victoria, Australia

## Abstract

Serum iron has long been thought to exhibit diurnal variation and is subsequently considered an unreliable biomarker of systemic iron status. Circadian regulation (endogenous ∼24-hour periodic oscillation of a biologic function) governs many critical physiologic processes. It is unknown whether serum iron levels are regulated by circadian machinery; likewise, the circadian nature of key players of iron homeostasis is unstudied. Here we show that serum iron, hepatic transferrin receptor (TFR1) gene (*Tfrc*) expression and erythropoietic activity exhibit circadian rhythms. Daily oscillations of serum iron, hepatic *Tfrc* expression and erythropoietic activity are maintained in mice housed in constant darkness, where oscillation reflects an endogenous circadian period. Oscillations of serum iron, hepatic *Tfrc* and erythropoietic activity were ablated when circadian machinery was disrupted in *Bmal1* knockout mice. Interestingly, we find that circadian oscillations of erythropoietic activity and hepatic *Tfrc* expression are maintained in opposing phase, likely allowing for optimised usage and storage of serum iron whilst maintaining adequate serum levels. This study provides the first confirmatory evidence that serum iron is circadian regulated and uncovers liver-specific circadian regulation of TFR1, a major player in cellular iron uptake.

## Introduction

Iron plays a crucial role in fundamental cellular processes throughout the body, including erythropoiesis. Iron is imperative for the transport of oxygen and as such, approximately 70% of bodily iron is locked in the erythroid compartment within hemoglobin. Red blood cells are the most abundant cells of the body with 200 billion new cells produced daily. Maintaining erythropoiesis at this rate requires 20mg of iron daily,^1^ underscoring the intricate connection between iron metabolism and erythropoiesis. Cellular iron requirements are supplied by plasma iron that circulates bound to transferrin. Circulating iron, typically measured clinically by assaying serum iron levels, has been recognised to exhibit diurnal variation over a 24-hour period since the 1940’s, with concentrations higher during the day and reduced during the night in humans.^2-4^ Partly for this reason, serum iron is generally considered an unreliable biomarker of iron status.^5,6^ Plasma iron levels are regulated by the cellular influx and efflux of iron. Iron entry to the plasma from the duodenum (dietary iron) or reticuloendothelial system (iron recycled from senescent red cells) occurs via the sole cellular iron export protein, ferroportin. Ferroportin function is governed by hepcidin, which is expressed by the liver in response to iron loading and inflammation and suppressed during iron deficiency and enhanced erythropoiesis.^1^ Transferrin receptor 1 (TFR1) facilitates uptake of iron from the plasma via receptor-mediated endocytosis of transferrin-bound iron. TFR1 is post-transcriptionally regulated by cellular iron status via binding of iron-regulatory proteins (IRPs) to iron response elements (IREs) in its 3’ region, allowing increased TFR1 expression when iron levels are low and reduced TFR1 expression when iron is replete.^7^

Circadian rhythms (endogenous ∼24-hour periodic oscillation of a biologic function) govern many critical physiologic processes, including hepatic metabolic systems.^8^ These physiological oscillations are synchronised by environmental signals (zeitgebers) such as light/dark cycles. Light sensed by the retina is connected to the suprachiasmatic nucleus (SCN) in the brain. The SCN, known as the master clock, is responsible for synchronizing peripheral clocks distributed across the organism. Food intake is also a potent zeitgeber and altered feeding can reset the phase of circadian genes in peripheral organs such as the liver.^9^ The circadian clock comprises the activating transcription factors CLOCK and its partner BMAL1 (gene: *ARNTL*), which activate transcription of the *PER* and *CRY* genes, whose protein products accumulate over the day and act to repress CLOCK-BMAL1 transcription, establishing oscillation. CLOCK-BMAL1 also activate expression of the repressive transcription factors REV-ERBα (gene: *NR1D1*) and REV-ERBβ,^10^ which compete with activating transcription factors RORα and RORγ; this establishes a second oscillation.^11^

Whether serum iron is regulated by endogenous circadian mechanisms, or simply reflects behavioural (e.g. dietary) factors, remains unknown. Furthermore, circadian oscillation of genes that govern iron homeostasis have not been defined. Here, we provide the first mechanistic evidence that clinically-observed oscillations in serum iron are governed by endogenous circadian mechanisms. Further, we show that hepatic TFR1 gene expression and erythropoietic activity also oscillate in a circadian fashion, in opposing phases.

## Results

We firstly determined if serum iron oscillates in wildtype C57Bl/6 mice maintained on a cycle of alternating 12-hours light and 12-hours darkness. After 2 weeks of entrainment, we collected samples every four hours from Zeitgeber time (ZT) 0 (lights turned on) through ZT12 (lights turned off) to ZT20. We initially confirmed the expected reciprocal oscillations in hepatic mRNA expression of the core circadian genes *Clock, Arntl, Nr1d1, Cry1 and Per2* (p<0.001; Figure 1A-E). In this system, serum iron levels oscillated, with a difference in mean serum iron of 10μmol/L from peak to trough (p<0.0001; Figure 1F). This variation in serum iron is unlikely to reflect changes in iron recycled by the reticuloendothelial system as splenic iron did not vary significantly across the day (p=0.995; Figure 1G).

**Figure 1.**
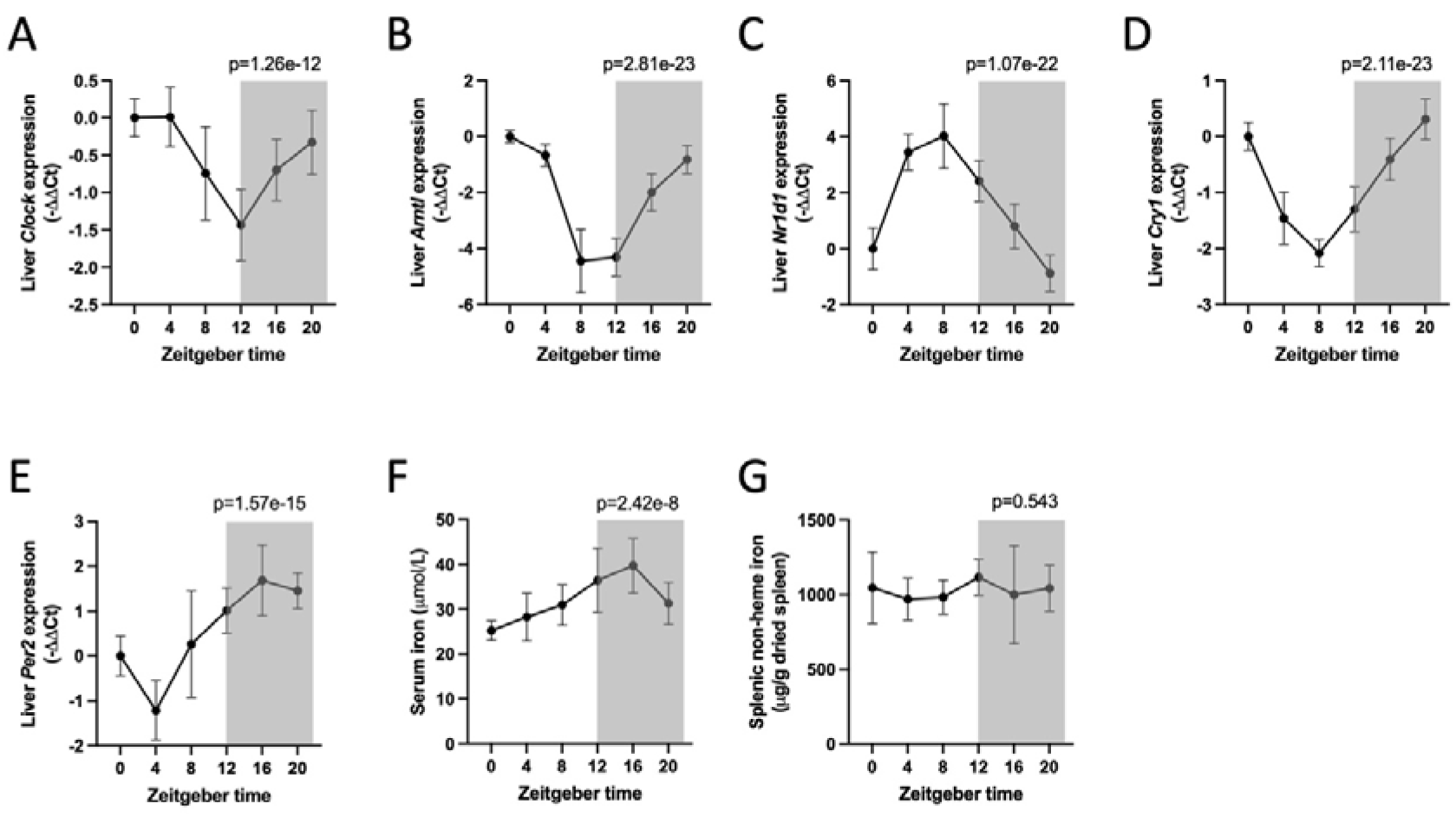
Expression of circadian genes, serum iron and splenic iron in mice maintained on a 12-hour light/ 12-hour dark cycle. C57Bl/6 mice were maintained on a 12-hour light / 12-hour dark cycle for two weeks before mice were euthanised and organs harvested every 4 hours. Liver *Clock* (A), *Arntl* (B), *Nr1d1* (C), *Cry1* (D) and *Per2* (E) expression (N=13 except ZT0 where N=11 and *Cry1* ZT8 where N=12). (F) Serum iron (N=13 except ZT0 and 8 where N=12) (G) Splenic non-heme iron (N=7 except ZT16 where N=5). P values derived from JTK_Cycle algorithm.

To confirm that oscillations in serum iron are not simply a reflection of timing of dietary iron intake, we performed time-restricted feeding experiments. Mice had feed restricted to the light (ZT0-ZT12) or dark hours (ZT12-ZT24). Mice fed in either condition consumed the same amount of food (Figure 2A). Night fed mice reflected *ad libitum* fed mice with liver *Clock* and *Arntl* expression reduced at ZT4 compared to ZT16, but increased liver *Per2* expression and serum iron levels (Figure 2B-E). In contrast, day feeding inversed the rhythm o f core circadian genes in the liver as previously reported^12^ (Figure 2B-D). However, day feeding did not lead to increased levels of serum iron in the daylight hours (ZT4) (Figure 2E), suggesting changes in serum iron levels are not solely a consequence of changes in dietary iron availability throughout the day.

**Figure 2.**
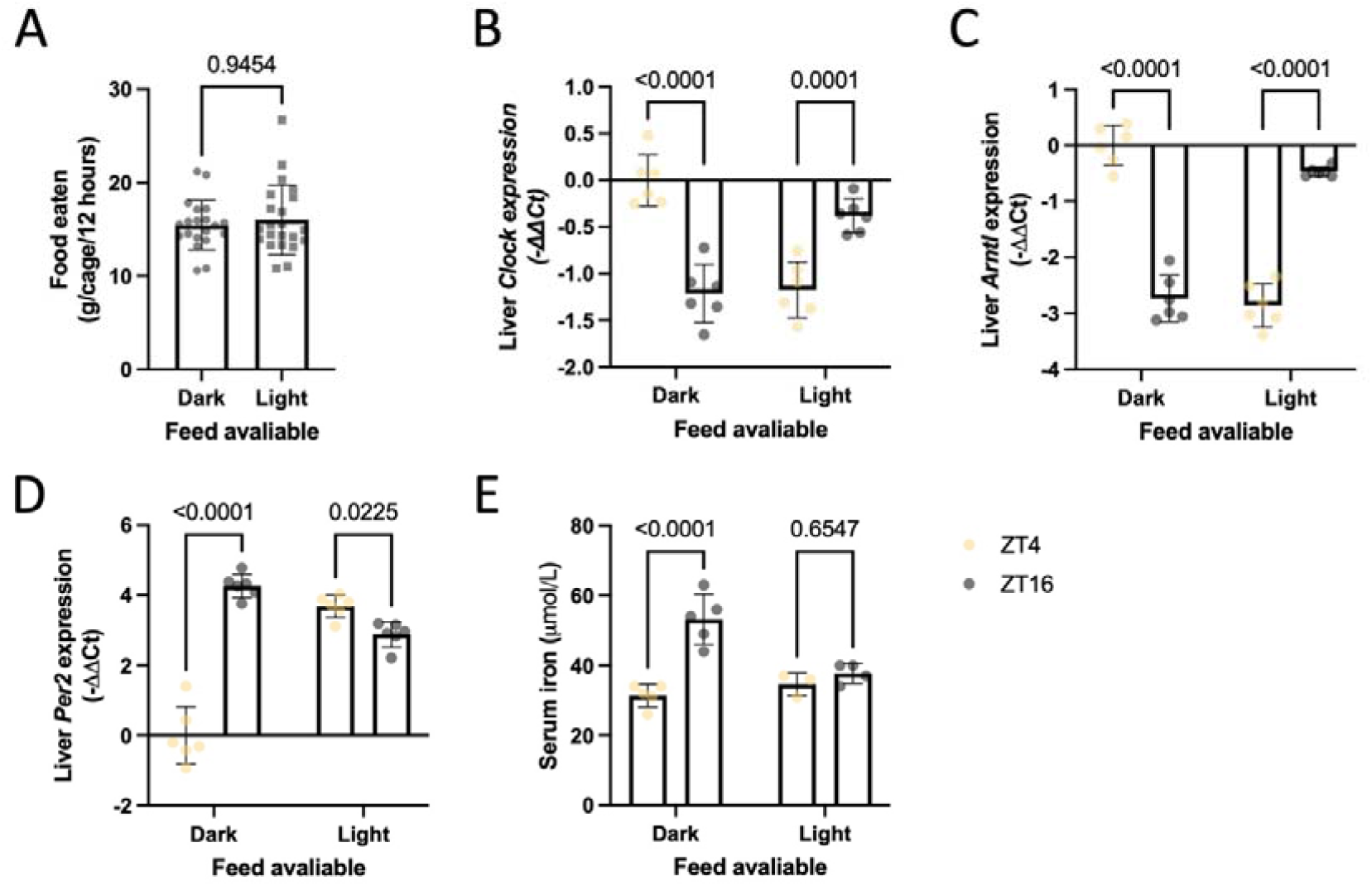
Expression of circadian genes and serum iron in mice with feed restricted to the light or dark hours. C57Bl/6 mice were maintained on a 12-hour light / 12-hour dark cycle with *ad libitum* access to food for two weeks before having feed restricted to the light hours - zeitgeber time (ZT)0 to ZT12 - or the dark hours - ZT12 to ZT0 - for 12 days before mice were euthanised and organs harvested at ZT4 or ZT16. (A) Food eaten per cage per 12 hours (N=20 dark fed, 22 light fed). Liver *Clock* (B), *Arntl* (C) and *Per2* (D) expression (N=6). (E) Serum iron (N=5 dark fed, 3 light fed ZT4 and 4 light fed ZT16). P values are derived from (A) Mann-Whitney test and (B-E) Two-way ANOVA with Šídák’s correction for multiple comparisons.

We initially hypothesised that variation in serum iron may be driven by changes in hepcidin but found that hepatic *Hamp1* mRNA was relatively stable across the day (adjusted (adj) p=0.225; Figure 3A) as was serum hepcidin (p=1.000; Figure 3B). We then hypothesised that other key players in iron regulation in the liver involved in either iron export – ferroportin (gene: *Slc40a1*) – or iron import – TFR1 (gene: *Tfrc*) – may have circadian oscillations that impact serum iron variation. We found that *Slc40a1* (adj p=0.763; Figure 3C) mRNA levels were relatively stable across the day. However, *Tfrc* mRNA expression exhibited variation (adj.p<0.0001, 1.9-fold change peak to trough; Figure 3D). Oscillations of *Tfrc* appear specific to the liver and were not observed in hematopoietic organs, i.e. bone marrow or spleen (Figure 3E-F). Interestingly, oscillations of *Tfrc* were in phase with oscillation of serum iron, suggesting hepatic iron uptake is increased when serum iron is abundant. We did not observe statistically significant rhythmic changes in liver iron content across the day in mice (p=0.068, Figure 3G). However, post-hoc analysis directly comparing liver non-heme iron at ZT16 (the timepoint at which both liver *Tfrc* expression and serum iron are at their zenith) to ZT4 (when liver *Tfrc* is at its nadir), confirmed higher levels of liver iron in mice at ZT16 (p=0.001). This finding is inconsistent with *Tfrc* expression being exclusively regulated by the IRE-IRP system, wherein *Tfrc* expression would be expected to be anti-phase to liver and/or serum iron levels.

**Figure 3.**
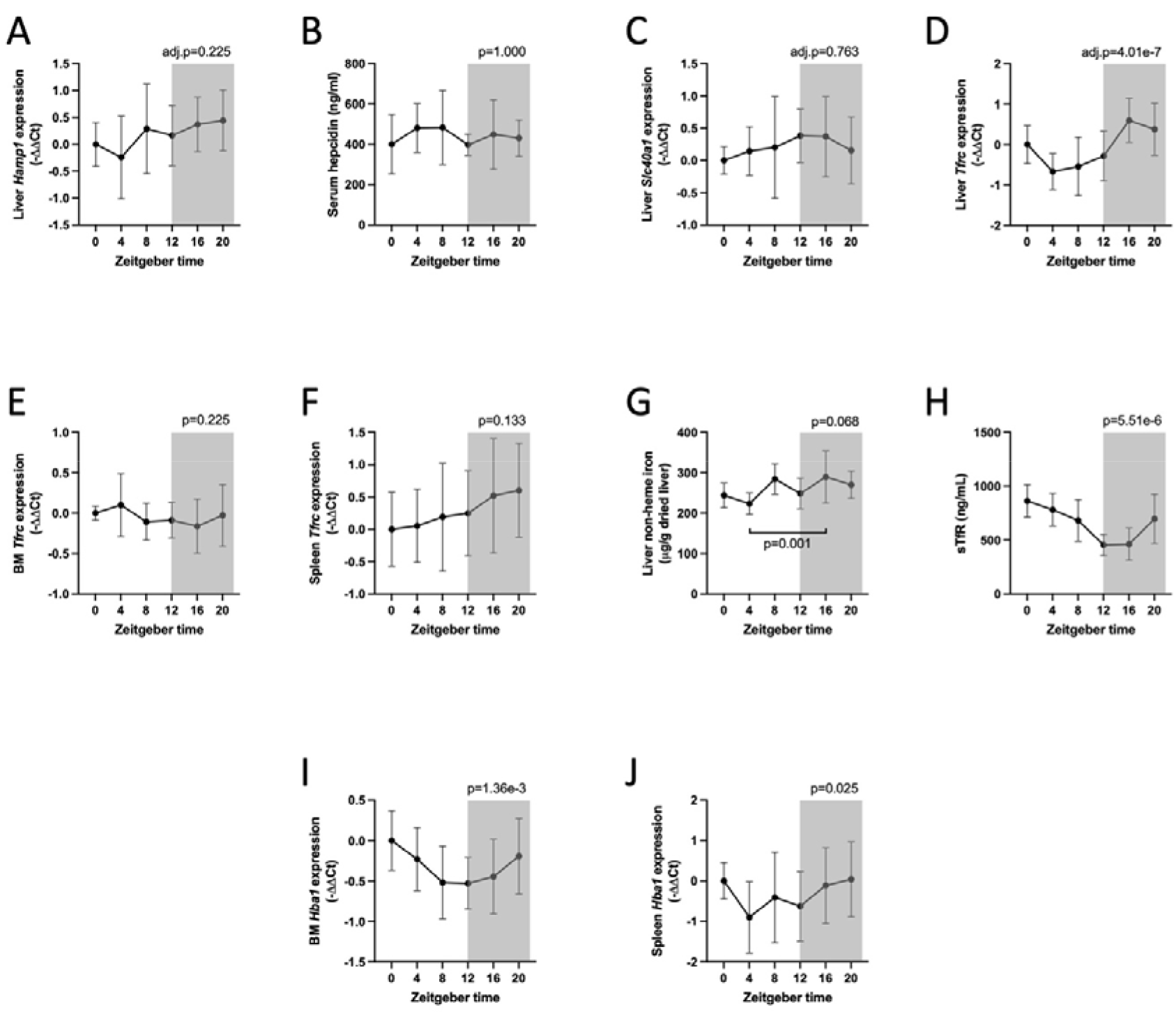
Changes in iron regulation and erythropoiesis in mice maintained on a 12-hour light/ 12-hour dark cycle. C57Bl/6 mice were maintained in constant darkness for 2 weeks before mice were euthanised and organs harvested every 4 hours. (A) Liver *Hamp1* expression (N=13 except ZT0 where N=11). (B) serum hepcidin (N=7 except ZT8 and 12 where N=6). Liver (C) *Scl40a1* and (D) *Tfrc* expression (N=13 except ZT0 where N=11). (E) Bone marrow (BM) *Tfrc* expression (N=12 except ZT4, 8 and 16 where N=13). (F) Spleen *Tfrc* expression (N=13 except ZT0 where N=12). (G) Liver non-heme iron (N=9 except ZT8 and 20 where N=10 and ZT0 where N=8). (H) Soluble transferrin receptor (sTfR) (N=7). (I) Bone marrow (BM) *Hba1* expression (N=12 except ZT4, 8 and 16 where N=13). (J) Spleen *Hba1* expression (N=13 except ZT0 where N=12). P values derived from JTK_Cycle algorithm. P value comparing ZT4 to ZT16 in (G) derived from Mann-Whitney test.

We next sought to confirm that *Tfrc* expression, but not hepcidin nor ferroportin gene expression, exhibits daily variation in human livers utilising the CircaDB database (http://circadb.hogeneschlab.org/human). CircaDB contains RNA-sequencing data of 13 human organs, including the liver, collected by the GTEx Consortium^13^ following CYCLOPS analysis^14^ which allowed reconstruction of temporal order in the absence of time-of-day information. This dataset suggests that hepatic *TFRC* (p=0.05) but not *HAMP* (p=0.66) nor *SLC40A1* (p=0.33) mRNA expression exhibits oscillation over a 24-hour period in humans.

We supposed that the observed oscillations in both hepatic *Tfrc* and liver iron levels incompletely explain the daily oscillations in serum iron and that iron utilisation in erythroid organs may change across the day, further contributing to the variation in serum iron. It has been reported that erythropoiesis (as expressed by changes in the percentage of reticulocytes) exhibits diurnal variation in both mice^15^ and humans^16^. In mice, reticulocyte levels peak around the time when lights are turned on (ZT0) and trough when lights are turned off (ZT12); i.e. in the opposite phase to serum iron. In keeping with this, serum soluble transferrin receptor (sTfR) concentration, a known marker of erythropoiesis^17^, showed variation with a difference in mean sTfR of 378ng/mL from peak to trough (p<0.0001; Figure 3H). Importantly, oscillations of sTfR were in the opposite phase to serum iron. To confirm diurnal changes in erythropoiesis, we looked at the expression *Hba1* in the bone marrow, the gene encoding alpha-globin. *Hba1* expression showed diurnal oscillations in phase with sTfR (p=0.001; Figure 3I). In mice, the spleen represents a significant site of erythropoiesis during infancy^18^ and in adulthood during stress erythropoiesis^19^. In young adult mice (8-to-9-week-old) under stress free conditions, diurnal oscillations of *Hba1* were also observed in the spleen (p=0.025; Figure 3J), reflecting residual steady state erythropoiesis in the spleen. These data indicate erythropoiesis exhibits diurnal variation.

To this point, we have purposefully described the observed oscillations as *diurnal* in nature. Next, we looked to discover whether the oscillations in serum iron, hepatic *Tfrc*, and erythropoiesis are endogenously *circadian* in nature and not simply occurring in response to external signals. To demonstrate thus, we undertook a free running experiment, in which animals were housed in constant darkness with *ad libitum* access to food and water for 2 weeks: conditions under which ongoing oscillation of parameters can be considered to reflect a true endogenous circadian period.^20^ Figure 4A-C demonstrates reciprocal hepatic oscillation of core clock genes *Clock, Arntl* and *Nr1d1*, as expected. In this free running context, we observed oscillation of serum iron (p<0.005; Figure 4D), with a difference in mean serum iron of 12μmol/L from peak to trough. Since hepcidin expression did not oscillate under 12-hour light / 12-hour dark conditions, we did not expect to observe significant oscillations in liver *Hamp1* expression in free running conditions. Indeed, *Hamp1* expression was stable across the day under complete darkness (p=1.000; Supplemental Figure 1A). In contrast, hepatic *Tfrc* expression appeared to oscillate under free running conditions, however this oscillation did not reach statistical significance when the JTK_Cycle period was set to 24 hours (p=0.123; Figure 4E). We hypothesised that this is likely due to the true endogenous period of hepatic *Tfrc* being greater than 24 hours. Indeed, post-hoc testing with the JTK_Cycle allowing a period of 32 hours revealed statistically significant oscillations in *Tfrc* expression (p<0.0001). We also observed oscillations of sTfR levels, although not quite reaching statistical significance (p=0.062; Figure 4F), with a mean difference of 212ng/mL from peak to trough. Again, lowest levels of sTfR aligned with the peak of hepatic *Tfrc* expression and serum iron levels.

**Figure 4.**
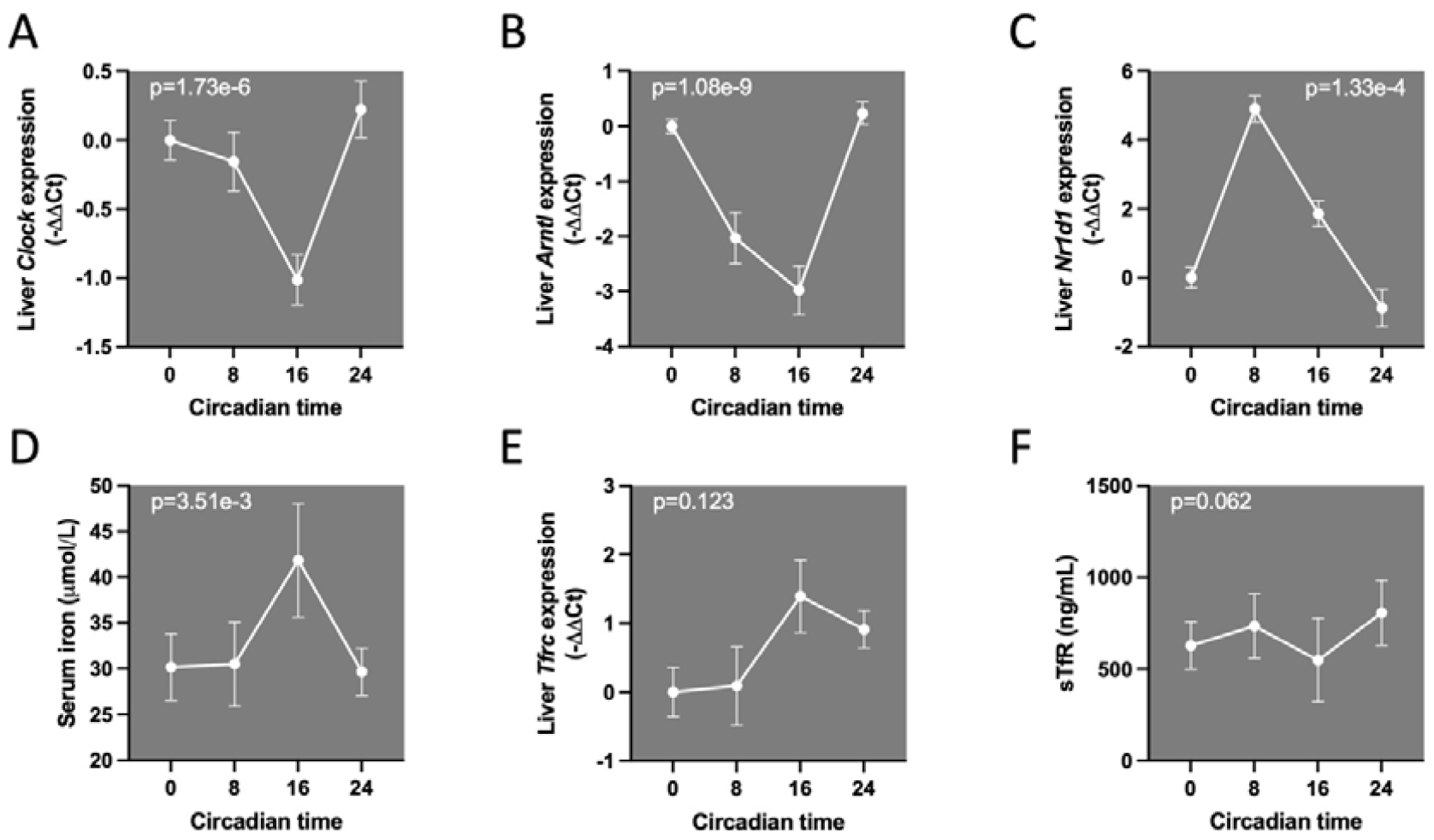
Expression of circadian and iron homeostatic genes, serum iron and soluble transferrin receptor in mice maintained in constant darkness. C57Bl/6 mice were maintained in constant darkness for 2 weeks before mice were euthanised and organs harvested. Liver *Clock* (A), *Arntl* (B) and *Nr1d1* (C) expression. (D) Serum iron. (E). Liver *Tfrc* expression. (F) Soluble transferrin receptor (sTfR). N=6. P values derived from JTK_Cycle algorithm.

Next, we assessed whether circadian genes influence oscillations of serum iron, hepatic *Tfrc* and erythropoiesis. We utilised a *Bmal1* knockout model,^21^ in a 12-hour light/12-hour dark setting. Littermate control (*Bmal1*^*+/+*^) mice were compared to *Bmal1* knockout (*Bmal1*^*-/-*^) animals at ZT4 and ZT16 (the timepoints at which we had initially observed the largest differential *Tfrc* expression). As expected, *Bmal1*^*+/+*^ mice exhibited oscillation of *Nr1d1* mRNA in the liver between these two timepoints, whereas *Nr1d1* expression was ablated at both timepoints in *Bmal1*^*-/-*^ animals (Figure 5A, genotype-time interaction p<0.0001), indicative of effective disruption of the circadian machinery. Whilst *Bmal1*^*+/+*^ mice exhibited an increase in serum iron between ZT4 and ZT16 as previously observed, this increase was ablated in *Bmal1*^*-/-*^ animals (Figure 5B, genotype-time interaction p=0.048). Genetic deletion of *Bmal1* had no impact on liver *Hamp1* as expected (Supplemental Figure 1B). In contrast, oscillations of hepatic *Tfrc* were ablated in *Bmal1*^*-/-*^ mice (p=0.795; Figure 5C); a finding that is supported by a published liver transcriptomic dataset of *Bmal1*^*-/-*^ and wildtype mice at ZT4 and ZT16.^22^ However, we were unable to confirm a statistically significant interaction between genotype and time for *Tfrc* expression. We then wondered whether systemic dysregulation of the circadian clock would have an impact on erythropoiesis. Interestingly, although sTfR levels were increased at ZT4 compared to ZT16 in *Bmal1*^*+/+*^ mice, levels were comparable at both timepoints in *Bmal1*^*-/-*^ animals (Figure 5D), suggesting circadian oscillations of erythropoiesis were impacted in these mice. In keeping with this, we observed no significant difference in the expression of the erythroid gene *Hba1* in the bone marrow of *Bmal1*^*-/-*^ animals at ZT4 compared to ZT16 (p=0.274; Supplemental Figure 2). These data imply a direct role for the circadian clock machinery in the bone marrow and liver in the regulation of serum iron via control of hepatic *Tfrc* expression and erythropoiesis.

**Figure 5.**
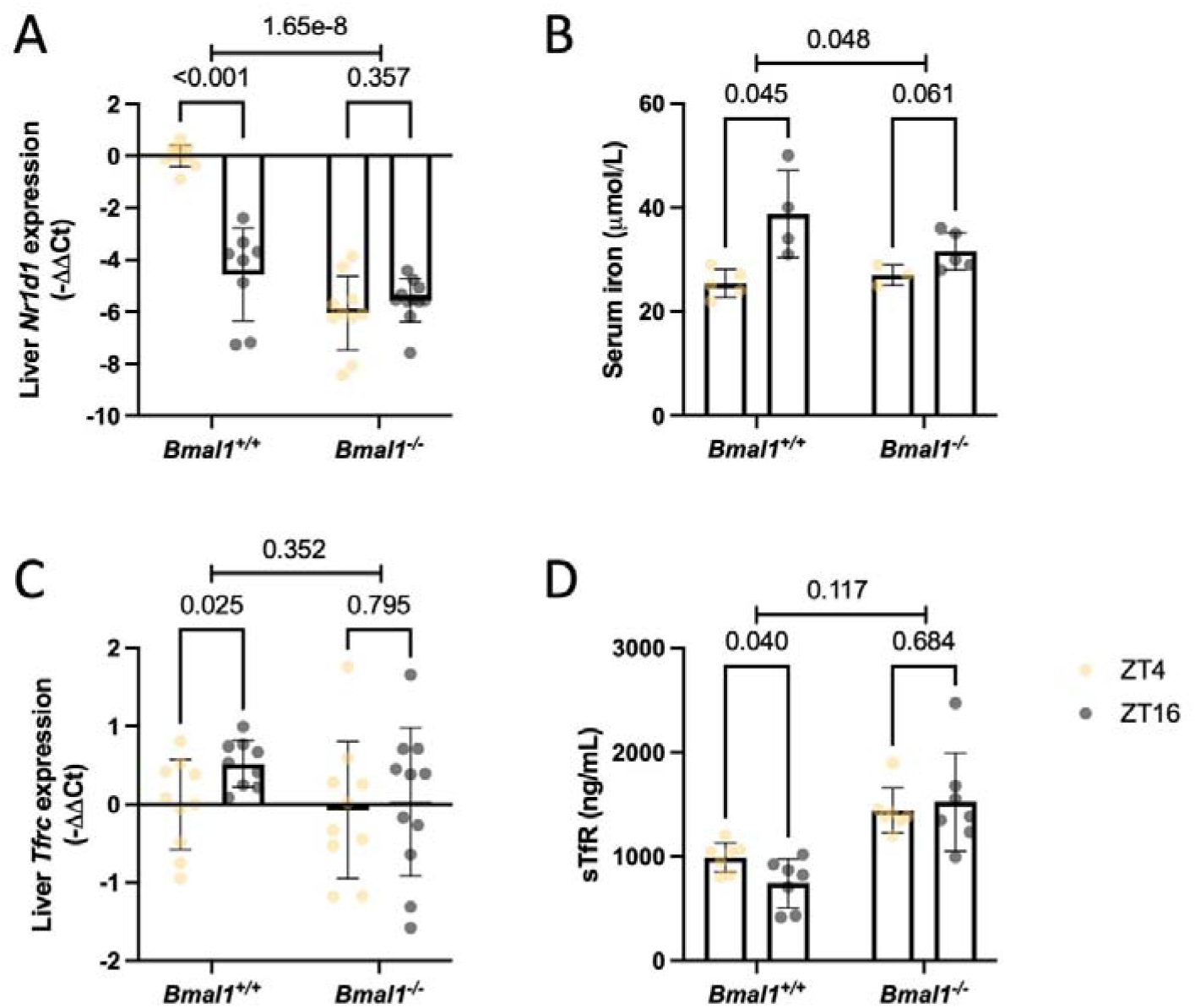
Expression of circadian and iron homeostatic genes, serum iron and soluble transferrin receptor in *Bmal1*^*-/-*^ mice. *Bmal1* knockout (*Bmal1*^-/-^) and littermate control (*Bmal1*^*+/*+^) mice were maintained a 12-hour light/12-hour dark cycle for 2 weeks before organs were harvested at zeitgeber time (ZT) 4 and ZT16. (A) Liver *Nr1d1* expression (N=10 *Bmal1*^*+/+*^ ZT4, 8 *Bmal1*^*+/+*^ ZT16, 10 *Bmal1*^*-/-*^ ZT4 and 11 *Bmal1*^*-/-*^ ZT16). (B) Serum iron (N=5 *Bmal1*^*+/+*^ ZT4, 4 *Bmal1*^*+/+*^ ZT16, 3 *Bmal1*^*-/-*^ ZT4 and 5 *Bmal1*^*-/-*^ ZT16). (C) Liver *Tfrc* expression (N=10 *Bmal1*^*+/+*^ ZT4, 9 *Bmal1*^*+/+*^ ZT16, 10 *Bmal1*^*-/-*^ ZT4 and 11 *Bmal1*^*-/-*^ ZT16). (D) Soluble transferrin receptor (sTfR; N=7). P values within genotypes are derived from Student’s T-test with Welch’s correction. P values across genotypes are derived from a likelihood ratio test of a genotype-timepoint interaction.

## Discussion

The mammalian circadian clock maintains rhythmic physiologic patterns in functions including sleep-wake cycles, feeding/fasting, and hormonal secretion, over a 24-hour period. Rhythms are synchronised across the whole organism via a central mechanism within the suprachiasmatic nucleus, and by peripheral clocks in tissues including the liver.^23^ Maintenance of oscillation during free running conditions, coupled with ablation of oscillation following genetic deletion of the afferent arm of the circadian clock, provide the first direct evidence of circadian oscillations of serum iron. This daily rhythmic oscillation appears to be a consequence of direct circadian regulation in two distinct iron regulatory organs, the bone marrow and the liver. Erythropoiesis in the bone marrow undergoes circadian oscillations as demonstrated here by oscillations of sTfR and *Hba1* expression. Erythropoiesis is the major consumer of bodily iron. Thus, it is not surprising that serum iron levels reach their nadir when erythropoiesis is at its peak, at the start of the nocturnal mouse’s resting phase. In humans, it has been shown that erythropoiesis is highest at 1am (i.e the resting phase),^16^ correlating with the lowest levels of serum iron.^2,4^ It is plausible that the endogenous circadian clock maintains erythropoiesis to the resting phase in mammals to ensure energy supply for this vital function.

On the other hand, the liver undergoes circadian oscillations of *Tfrc* expression in the opposite phase to erythropoiesis. We hypothesise that the mammalian circadian regulation has evolved in this way to optimise storage of excess iron from the serum when it is not required for erythropoiesis. In keeping with this, we observed higher levels of liver iron at times when erythropoiesis was low (ZT16) compared to times when erythropoiesis was high (ZT4). Transferrin receptor 1 is post-transcriptionally regulated via the IRE-IRP system which stabilises *Tfrc* mRNA in conditions of cellular iron deficiency (to enable further iron uptake). However, under basal conditions, circadian increases in hepatic *Tfrc* aligned with timepoints exhibiting higher serum and liver iron levels, suggesting circadian regulation of *Tfrc* is distinct from the IRE-IRP machinery. Further, we did not observe daily variation of other hepatic IRP-IRE controlled genes under 12-hour light/12-hour dark conditions – 3’ IRE, *Slc11a2* (Supplemental Figure 3A); 5’ IRE, *Slc40a1* (Figure 3C) and *Fth1* (Supplemental Figure 3B).

Interestingly, we found little evidence of diurnal oscillations of hepcidin, the “master” regulator of bodily iron, in mice. In keeping with this, the CircaDB database does not predict hepcidin gene expression to show diurnal variation in human liver (P=0.66). Previous clinical evidence has indicated evidence of diurnal variation of serum hepcidin in humans, in antiphase to serum iron.^24-26^ Levels of serum hepcidin in humans is clearly influenced by feed/fasting. Prolonged fasting led to 3.5-fold increase in serum hepcidin, whereas unrestricted caloric intake post-fasting significantly reduced serum hepcidin to levels 2-fold lower than baseline.^25^ These changes in hepcidin in response to feed/fasting in humans have been hypothesised to maintain serum iron levels in line with erythropoiesis in the fed/fasted state. In line with this, we observed a significant increase in hepcidin mRNA expression in the liver of mice who had feed restricted to the dark hours and therefore underwent a daily 12 hour fast (Supplemental Figure 4), whereas we saw no change in hepcidin levels in *ad libitum* fed mice (Figure 3A-B). These data indicate hepcidin is not circadian-regulated but instead is regulated by feeding/fasting through other mechanisms.

Here, we demonstrate evidence for factors driving the circadian variation of serum iron. We show a circadian network spanning two major iron regulatory organs, the bone marrow and liver, that work in harmony to maintain serum iron levels. Erythropoiesis depletes serum iron levels in the resting phase, whereas hepatic *Tfrc* expression is increased in the active phase optimising storage of excess serum iron when erythropoiesis is low. This mechanism is visually represented in Figure 6. Organ-specific oscillation of hepatic, but not haematopoietic *Tfrc* expression, potentially reflect differing roles of iron uptake between these organs. Our data provide the first evidence that long recognised oscillations in serum iron are indeed driven by endogenous circadian machinery, potentially linked to variations in erythropoiesis and liver iron uptake across the day.

**Figure 6.**
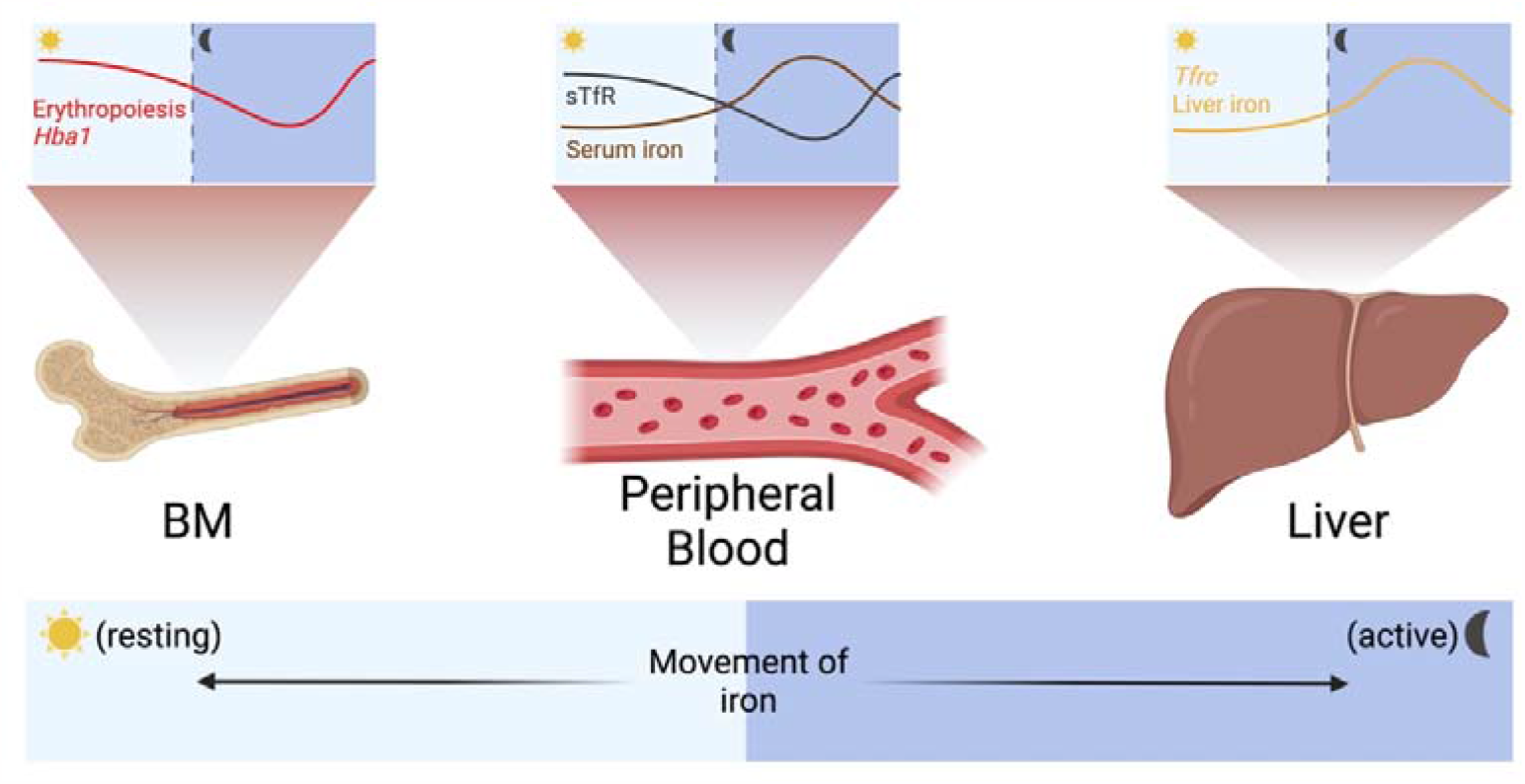
Serum iron is circadian regulated and linked to circadian networks across the liver and bone marrow working in harmony. Erythropoiesis, and thus *Hba1* expression, in the bone marrow (BM) is high during the daylight hours (resting phase), causing high levels of soluble transferrin receptor (sTfR) in the peripheral blood. This increased erythropoietic activity depletes serum iron levels in the resting phase. During the resting phase liver iron stores and hepatic *Tfrc* expression reach their nadir. However, during the dark hours (active phase), erythropoiesis levels decline, and serum iron levels increase. Harmoniously, expression of hepatic *Tfrc* peaks, allowing increased liver iron storage when serum iron levels are abundant. Created with BioRender.com.

## Methods

### Animals and Ethics

Mice were caged in pathogen free conditions with food (containing 180 mg/kg iron) and water available *ad libitum*. Lights were turned on at 6am (ZT0) and turned off at 6pm (ZT12). For time-restricted feeding experiments, feed was available from ZT0-ZT12 for light fed mice and ZT12-ZT0 for dark fed mice for 12 days; water was available *ad libitum*. For free running experiments, mice were housed in a temperature and humidity controlled dark box that blocked all external light. *Bmal1*^*-/-*^ mice^21^ were acquired from The Jackson Laboratory (strain: 009100) and bred inhouse. For all other experiments, wildtype C57Bl/6 mice were used. Male mice were used with age matched controls. The use of mice was in accordance with requirements set out by WEHI Animal Ethics Committee (approvals 2017.031 and 2020.034).

### Gene expression analysis

RT-qPCR from liver, spleen and bone marrow samples was performed as previously described.^27^ Data is presented as -ΔΔCt or -ΔCt, where ΔCt = Ct _target gene_ – Ct _*Hprt1*_ and ΔΔCt = ΔCt _sample_ – mean ΔCt _control (ZT0; ZT4 dark fed; CT0 or *Bmal1+/+* ZT4)_. Primers used for RT-qPCR are in the table below:

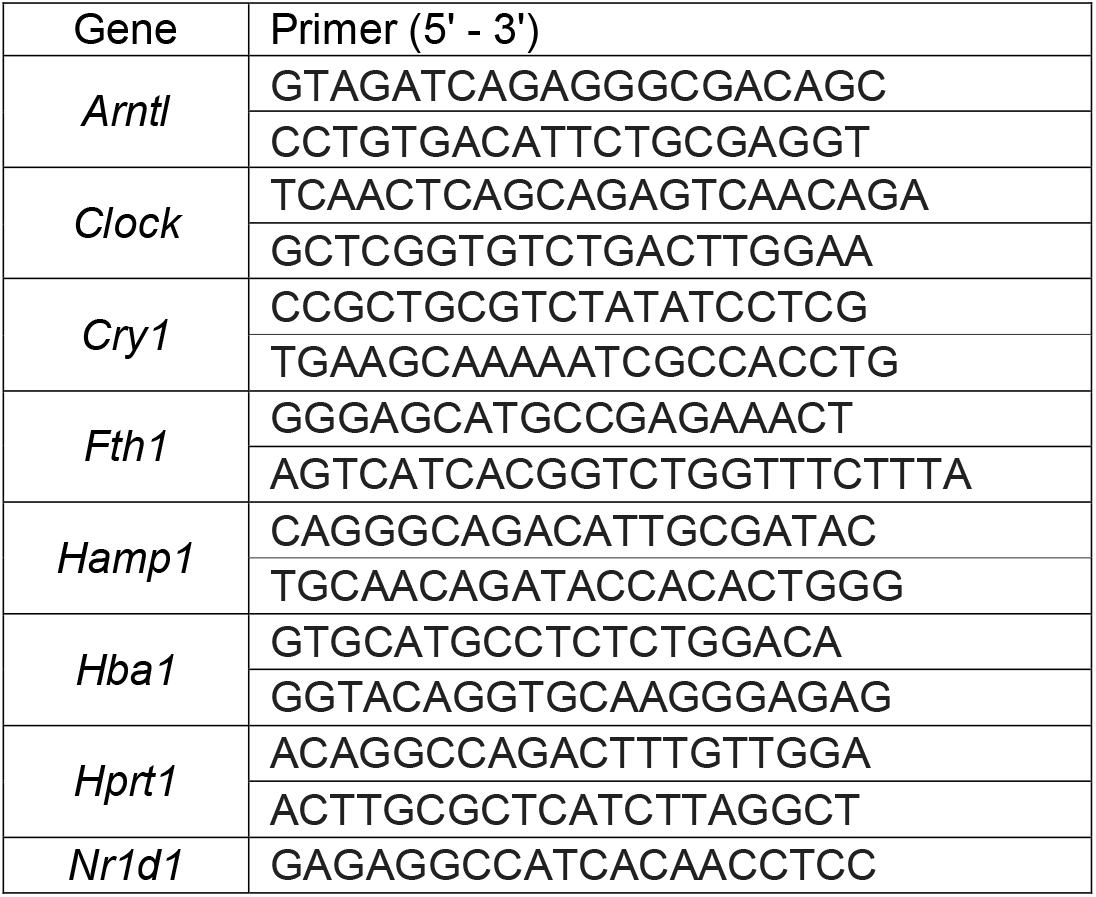
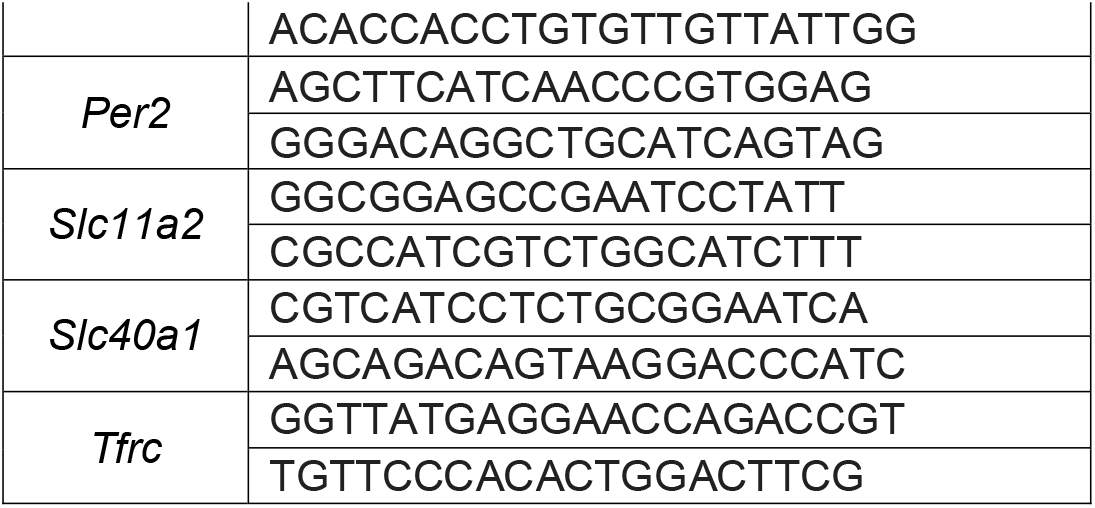

### Analysis of iron parameters

Serum iron and liver and spleen non-heme iron was measured as previously described.^27^ Serum sTfR and hepcidin were measured by ELISA (Abcam and Intrinsic Life Sciences, respectively).

### Statistical analysis

Biological functions were tested for the presence of a circadian oscillation with a period of 24 hours using the JTK_Cycle alrgorithm^28^ or post-hoc allowing a period of 24 or 32 hours. For the iron transporters (*Tfrc, Hamp1* and *Slc40a1*) in C57Bl/6 mice under 12-hour light/ 12-hour dark conditions, we report a bonferonni adjusted p-value. For spleen iron, assay 2 outputs were adjusted for an inter-assay batch effect by multiplying the original value by an adjustment factor calculated by taking the average relative difference (mean assay 1/mean assay 2) for each ZT timepoint where at least two samples had been run in both batches. Differences at ZT4 vs ZT16 in animals with feed restricted to the light or dark hours were compared by Two-way ANOVA with Šídák’s correction for multiple comparisons. Differences at ZT4 vs ZT16 in *Bmal1*^*+/+*^ and *Bmal1*^*-/-*^ mice were compared by Student’s T-test with Welch’s correction. A test for a timepoint by genotype interaction was conducted with a likelihood ratio test comparing a two-way ANOVA with an interaction term to a model without the interaction term. Sample size is denoted in the figure legends. Error bars represents mean±standard deviation. Statistical testing was performed in Prism 9.3.1; GraphPad Software and Stata v17.0.

## Acknowledgments

The authors thank the WEHI Bioservices facility for husbandry of all animals and assistance with animal experiments.

This work was supported by the National Health and Medical Research Council (GNT1159171, GNT1158696, GNT2009047 and GNT1123000 to S-R.P.; GNT1194345 to M.E.B.; and 2008090 to C.S). The contents of this published material are solely the responsibility of the individual authors and do not reflect the views of the NHMRC or funding partners. This work was also made possible through the Victorian State Government Operational Infrastructure Support and Australian Government National Health and Medical Research Council (NHMRC) Independent Research Institute Infrastructure Support Scheme (IRIISS).

## Author Contributions

Conceptualized study – C.B., and S-R.P.; Designed research – C.B., M.E.B., C.S., and S-R.P.; Performed research – C.B., A.P. and C.S.; Analyzed data – C.B., A.P., A.R.D.M, R.H.; Wrote the manuscript – C.B., A.P., A.R.D.M, R.H., M.E.B., C.S., and S-R.P.

