## Supplemental Data for "Serum iron variation is circadian-regulated and linked to the harmonic circadian oscillations of erythropoiesis and hepatic *Tfrc* expression in mice"

**Figures**

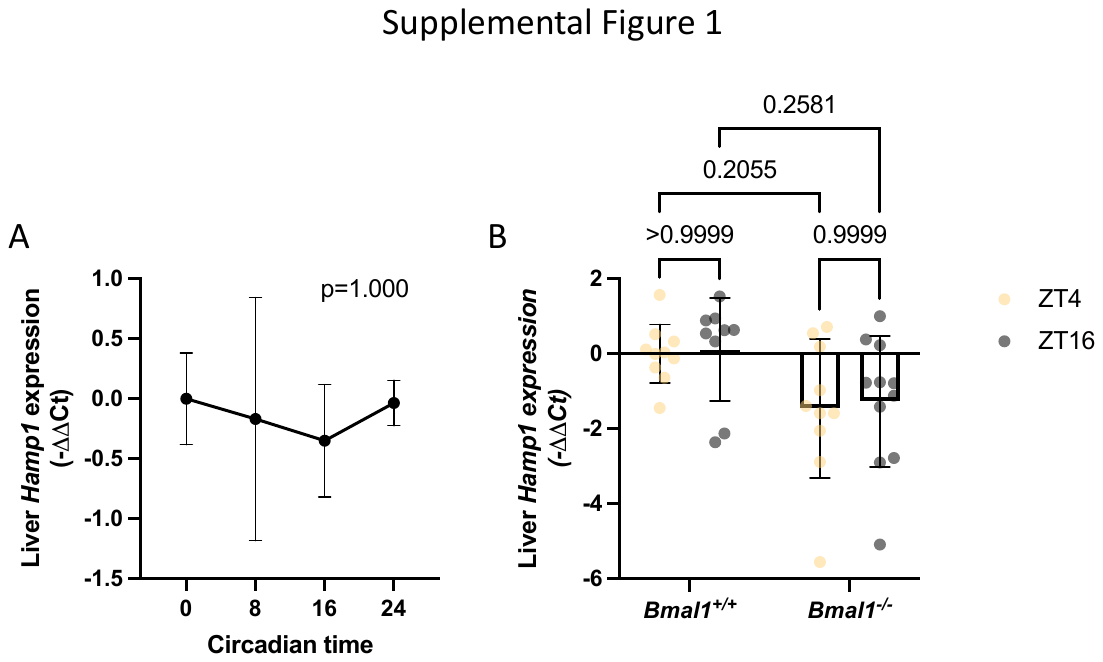

**Supplemental Figure 1 – Expression of hepcidin mRNA in mice maintained in constant darkness and in *Bmal1^-/-^* mice.** (A) C57Bl/6 mice were maintained in constant darkness for 2 weeks before mice were euthanised and organs harvested. Liver *Hamp1* expression (N=6; P values derived from JTK_Cycle algorithm). (B) *Bmal1* knockout (*Bmal1*^-/-^) and littermate control (*Bmal1^+/^*^+^) mice were maintained on a 12-hour light/12-hour dark cycle for 2 weeks before organs were harvested at zeitgeber time (ZT) 4 and ZT16. Liver *Hamp1* expression (N=10 *Bmal1^+/+^* ZT4, 8 *Bmal1^+/+^* ZT16, 10 *Bmal1^-/-^* ZT4 and 11 *Bmal1^-/-^* ZT16; P values derived from Two-way ANOVA with Šídák’s correction for multiple comparisons).

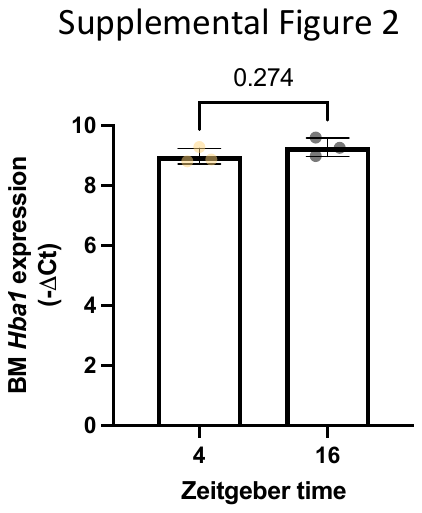

**Supplemental Figure 2 – Expression of *Hba1* in *Bmal1^-/-^* mice.** *Bmal1* knockout (*Bmal1*^-/-^) mice were maintained on a 12-hour light/12-hour dark cycle for 2 weeks before organs were harvested at zeitgeber time (ZT) 4 and ZT16. Bone marrow (BM) *Hamp1* expression (N=3; P values derived Student’s T-test with Welch’s correction).

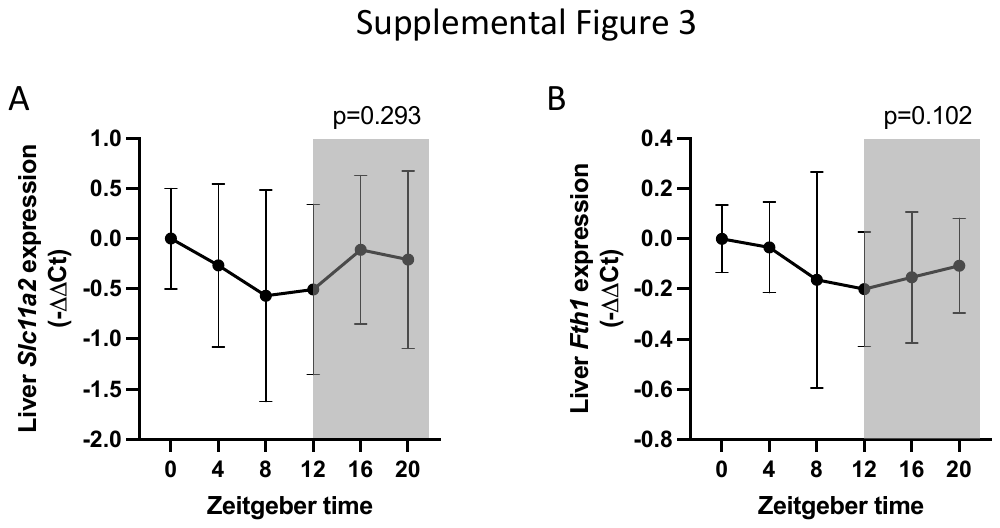

**Supplemental Figure 3 – Changes in IRE-IRP controlled genes in mice maintained on a 12-hour light/ 12-hour dark cycle.** C57Bl/6 mice were maintained in constant darkness for 2 weeks before mice were euthanised and organs harvested every 4 hours. Liver (A) *Slc11a2* and (B) *Fth1* expression (N=13 except ZT0, where N=11; P values derived from JTK_Cycle algorithm).

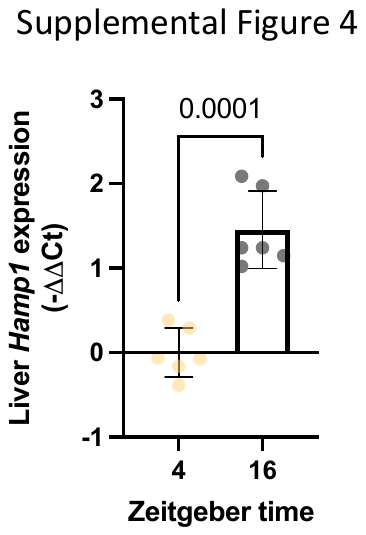

**Supplemental Figure 4 – Expression of hepcidin mRNA in mice with feed restricted to the dark hours.** C57Bl/6 mice were maintained on a 12-hour light / 12-hour dark cycle with *ad libitum* access to food for two weeks before having feed restricted to the dark hours - ZT12 to ZT0 - for 12 days before mice were euthanised and organs harvested at ZT4 or ZT16. Liver *Hamp1* expression (N=6). P values derived from Student’s T-test with Welch’s correction.
